# EffectorP 3.0: prediction of apoplastic and cytoplasmic effectors in fungi and oomycetes

**DOI:** 10.1101/2021.07.28.454080

**Authors:** Jana Sperschneider, Peter N. Dodds

**Affiliations:** Biological Data Science Institute, The Australian National University, Canberra, Australia; Black Mountain Science and Innovation Park, CSIRO Agriculture and Food, Canberra, Australia

## Abstract

Many fungi and oomycete species are devasting plant pathogens. These eukaryotic filamentous pathogens secrete effector proteins to facilitate plant infection. Fungi and oomycete pathogens have diverse infection strategies and their effectors generally do not share sequence homology. However, they occupy similar host environments, either the plant apoplast or plant cytoplasm, and may therefore share some unifying properties based on the requirements of these host compartments. Here we exploit these biological signals and present the first classifier (EffectorP 3.0) that uses two machine learning models: one trained on apoplastic effectors and one trained on cytoplasmic effectors. EffectorP 3.0 accurately predicts known apoplastic and cytoplasmic effectors in fungal and oomycete secretomes with low estimated false positive rates of 3% and 8%, respectively. Cytoplasmic effectors have a higher proportion of positively charged amino acids, whereas apoplastic effectors are enriched for cysteine residues. The combination of fungal and oomycete effectors in training leads to a higher number of predicted cytoplasmic effectors in biotrophic fungi. EffectorP 3.0 expands predicted effector repertoires beyond small, cysteine-rich secreted proteins in fungi and RxLR-motif containing secreted proteins in oomycetes. We show that signal peptide prediction is essential for accurate effector prediction, as EffectorP 3.0 recognizes a cytoplasmic signal also in intracellular, non-secreted proteins. EffectorP 3.0 is available at http://effectorp.csiro.au.

## Introduction

Eukaryotic filamentous pathogens encompass fungi and oomycetes that cause extensive damage to plants, including those providing food, fibre and biofuels (Savary *et al.*, 2019). Whilst oomycetes and fungi are not close phylogenetic relatives, they have very similar filamentous morphology and infection structures. These pathogens can broadly be classified according to their infection strategy: biotrophic pathogens grow and feed on living host tissue, hemibiotrophic pathogens switch to a necrotrophic phase after feeding on living tissues for some time and necrotrophic pathogens kill host cells and feed on dead tissues (Glazebrook, 2005). Pathogens and symbionts of plants use secreted proteins termed effectors to facilitate infection by suppressing plant defense responses and altering host cell structure and function (Lo Presti *et al.*, 2015). Pathogens secrete effectors into the host extracellular space, the plant apoplast, as well as into the intracellular space, the plant cytoplasm. Apoplastic effectors can either function in the apoplast (De Wit, 2016) or they can be bound to the fungal cell wall (Tanaka & Kahmann, 2021). In contrast, cytoplasmic effectors are delivered into the plant cell and some may subsequently target specific plant cell compartments. Effector delivery into host cells is well-characterized in bacteria, nematodes and insects (Mitchum *et al.*, 2013; Rodriguez & Bos, 2013; Galan *et al.*, 2014). For example, bacteria have conserved secretion systems such as the Type III Secretion System (T3SS) to directly inject effectors into the plant cell and nematodes use a stylet to infect effectors into the host cell.

Effector delivery mechanisms in fungi and oomycetes remains elusive (Lo Presti & Kahmann, 2017). In *Magnaporthe oryzae*, cytoplasmic effectors preferentially accumulate in the biotrophic interfacial complex (BIC) (Khang *et al.*, 2010) whereas apoplastic effectors follow the conventional secretory pathway (Giraldo *et al.*, 2013). In *Ustilago maydis*, a stable protein complex of five effectors and two membrane proteins has been suggested to be involved in effector translocation, however these proteins are not broadly conserved across fungi (Ludwig *et al.*, 2021). The presence of conserved sequence motifs in sequence-divergent oomycete effectors has been implicated in effector delivery (Whisson *et al.*, 2007). The RxLR motif occurs in the N-terminal sequence after the signal peptide cleavage site and is often followed by an acidic stretch of amino acids. However, experimental validation of RxLR-mediated effector uptake has been debated (Ellis & Dodds, 2011), with recent evidence pointing towards a role of the RxLR motif in effector cleavage before secretion (Wawra *et al.*, 2017). Furthermore, the RxLR motif is not conserved outside *Phytophthora* species and downy mildews. The WY domain is a structural domain associated with cytoplasmic oomycete effectors from the *Peronosporales* that has been used to identify effector candidates with degenerate RxLR motifs (Wood *et al.*, 2020). Similarly, degenerate RxLR motif searches have been applied in *Pythium* species that lack effectors with a canonical RxLR motif (Ai *et al.*, 2020). The LxLFLAK sequence motif defines the crinkling- and necrosis-inducing family (CRN) proteins and is also commonly used to search for CRN effectors (Schornack *et al.*, 2010). Whilst in oomycetes sequence motif and Hidden Markov Model searches are well-established methods to predict some classes of cytoplasmic effectors, it will miss *bona fide* effectors that do not carry such motifs or domains as well as apoplastic effectors (Wood *et al.*, 2020).

Computational effector prediction in fungi is challenging as fungal effectors do not share sequence similarities or conserved sequence motifs. In fungi, early approaches have been biased towards prioritizing small, cysteine-rich proteins as effector candidates (Stergiopoulos & de Wit, 2009; Sperschneider *et al.*, 2015a). More recently, machine learning methods have been applied to fungal effector prediction and their localization (Jones *et al.*, 2018). However, the current version of EffectorP has been trained only on fungal effectors, not oomycete effectors, and does not return the most likely localization of the effector (Sperschneider *et al.*, 2016; Sperschneider *et al.*, 2018a). Whilst ApoplastP predicts localization of plant and pathogen proteins to the plant apoplast (Sperschneider *et al.*, 2018b), it has not been trained to distinguish effector proteins from non-effector proteins. A deep learning method called deepredeff can predict effectors in bacteria, fungi and oomycetes (Kristianingsih & MacLean, 2021), but does not return effector localization. EffectorO is a machine learning method for effector prediction trained on the N-terminus of effector sequences, but it is applicable to oomycetes only (Nur *et al.*, 2021). Here, we introduce EffectorP 3.0, the first method that can predict if a secreted fungal or oomycete protein is one of these three types: an apoplastic effector, a cytoplasmic effector or a non-effector.

## Results

### Training of EffectorP 3.0 for prediction of apoplastic and cytoplasmic effectors

We first curated positive training sets of fungal and oomycete effectors with experimental support. We classified these effectors into apoplastic or cytoplasmic according to experimental evidence from the literature. If such evidence is absent, we classified effectors according to the infection strategy of the pathogen. After sequence homology reduction, 64 apoplastic effectors (50 from fungi, 14 from oomycetes) and 112 cytoplasmic effectors (77 from fungi, 35 from oomycetes) remained as positive training data (Table 1). For the negative training sets, we predicted secretomes from a wide range of fungal and oomycete genomes (Supplementary Table S1). We curated five secreted protein sets that are likely depleted in different effector classes: 1) a homology-reduced set of secreted proteins with an RxLR motif from plant-pathogenic oomycetes together with secreted proteins from biotrophic or hemibiotrophic fungi (*n* = 25,643; likely depleted in apoplastic effectors); 2) a homology-reduced set of secreted proteins without an RxLR motif from plant-pathogenic oomycetes together with secreted proteins from necrotrophic or hemibiotrophic fungi (*n* = 30,912; likely depleted in cytoplasmic effectors); 3) a set of secreted fungal saprophyte proteins and secreted proteins from brown algae (*n* = 28,080; likely depleted in effectors); 4) a set of secreted fungal ectomycorrhizal proteins (*n* = 14,357; likely depleted in cytoplasmic effectors); and 5) a set of secreted proteins from animal-pathogenic fungi and animal-pathogenic oomycetes (*n* = 5,181; likely depleted in apoplastic effectors). Proteins in the negative sets that share sequence homology to an effector in the positive training set were excluded.

**Table 1:**
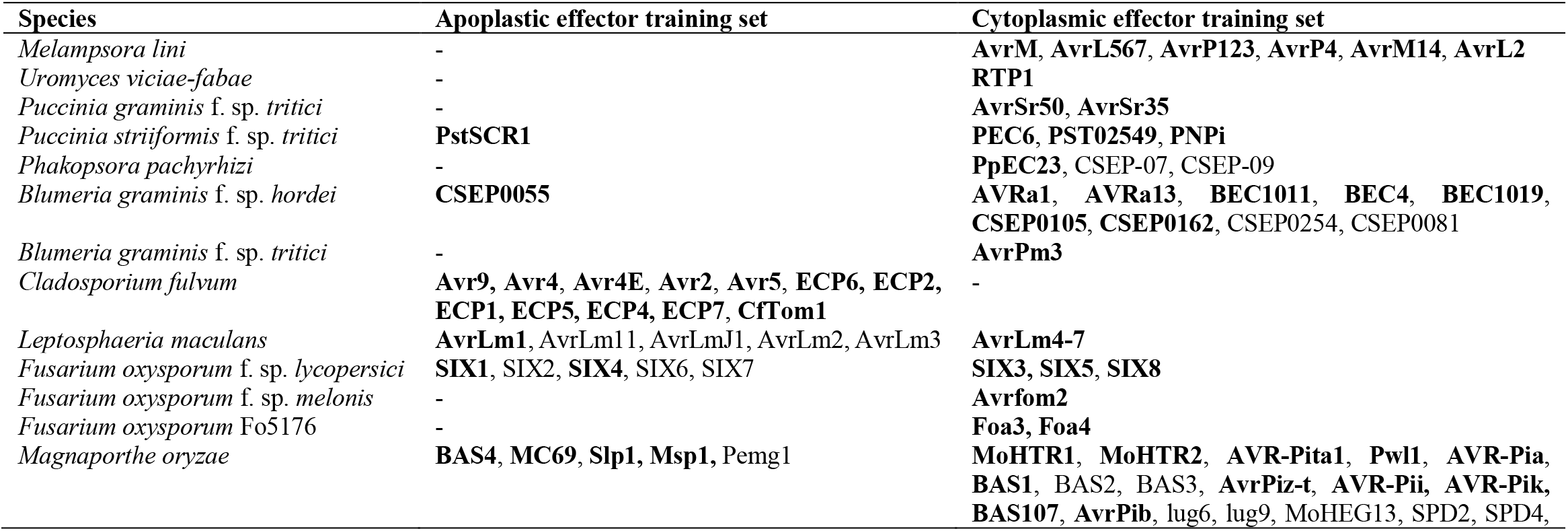
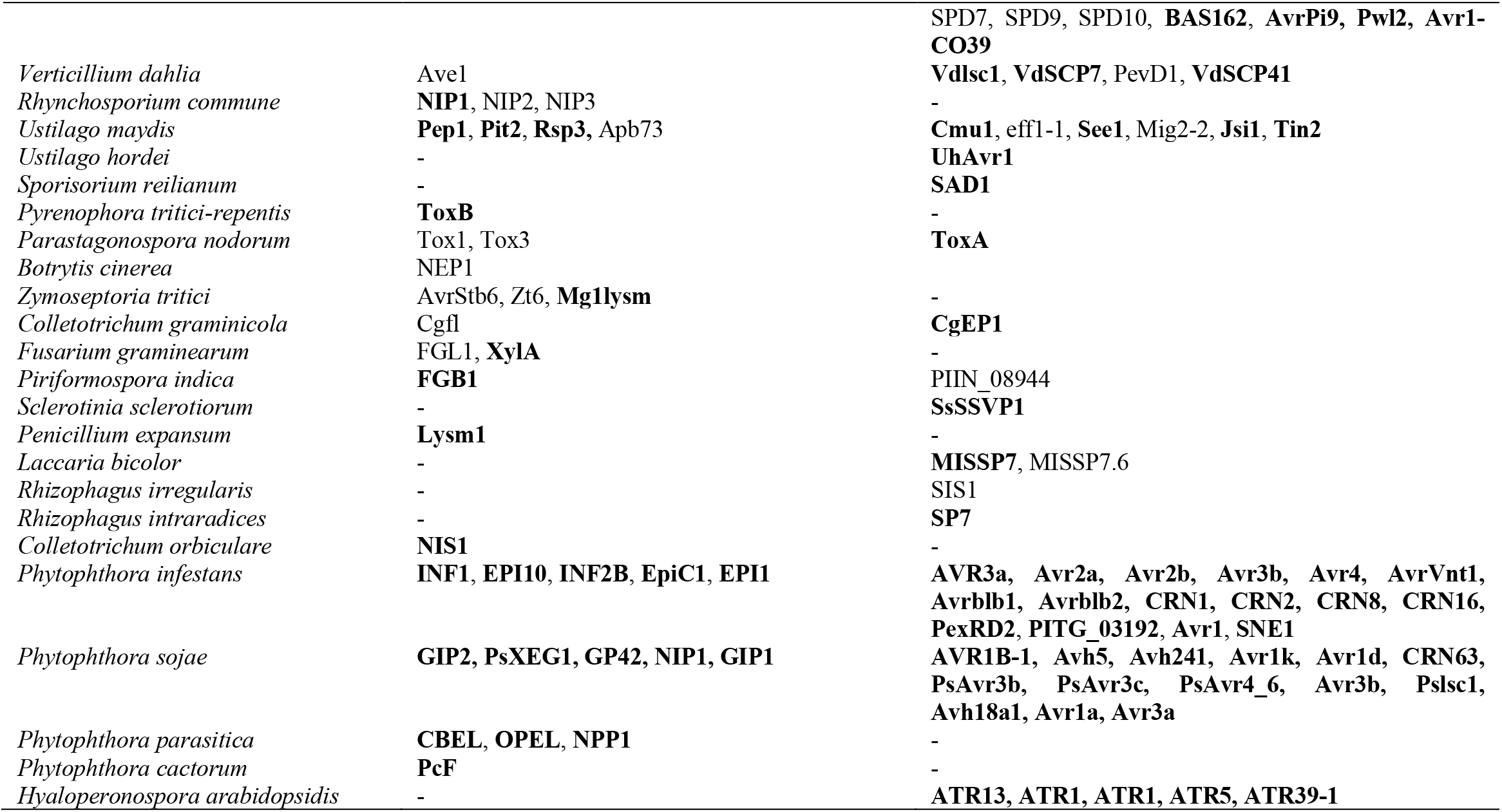
Fungal and oomycete effectors that were used in the training of EffectorP 3.0. Effectors with literature evidence for apoplastic or cytoplasmic localization are highlighted in bold.

We used an ensemble learning approach of classifiers that each are trained on a different subset of negative training data (Figure 1). Overall, we chose a total of 60 best‐performing models (Supplementary Table S2) comprising: Five Naïve Bayes classifiers and five C4.5 decision trees that discriminate between apoplastic effectors and secreted pathogen proteins; five Naïve Bayes classifiers and five C4.5 decision trees that discriminate between apoplastic effectors and secreted saprophyte/brown algae proteins; five Naïve Bayes classifiers and five C4.5 decision trees that discriminate between apoplastic effectors and secreted animal-pathogenic pathogen proteins; five Naïve Bayes classifiers and five C4.5 decision trees that discriminate between cytoplasmic effectors and secreted pathogen proteins; five Naïve Bayes classifiers and five C4.5 decision trees that discriminate between cytoplasmic effectors and secreted saprophyte/brown algae proteins; five Naïve Bayes classifiers and five C4.5 decision trees that discriminate between cytoplasmic effectors and secreted fungal ectomycorrhizal proteins. To generate EffectorP 3.0, we combined these 30 apoplastic and 30 cytoplasmic models into an ensemble classifier where each model has seen a different subset of negative training data and, for a given protein sequence input, returns a probability of whether it is 1) an apoplastic effector; 2) a cytoplasmic effector or 3) a non‐effector. If a secreted protein is predicted as both an apoplastic and cytoplasmic effector, EffectorP 3.0 reports it as dual-localized and the class with the highest probability as its primary localization. As the number of cytoplasmic fungal effectors is sufficiently large, we also used this positive set to train a fungal-specific cytoplasmic model with the same negative data sets and then combined this model with the apoplastic model as above to generate EffectorP-fungi 3.0.

**Figure 1:**
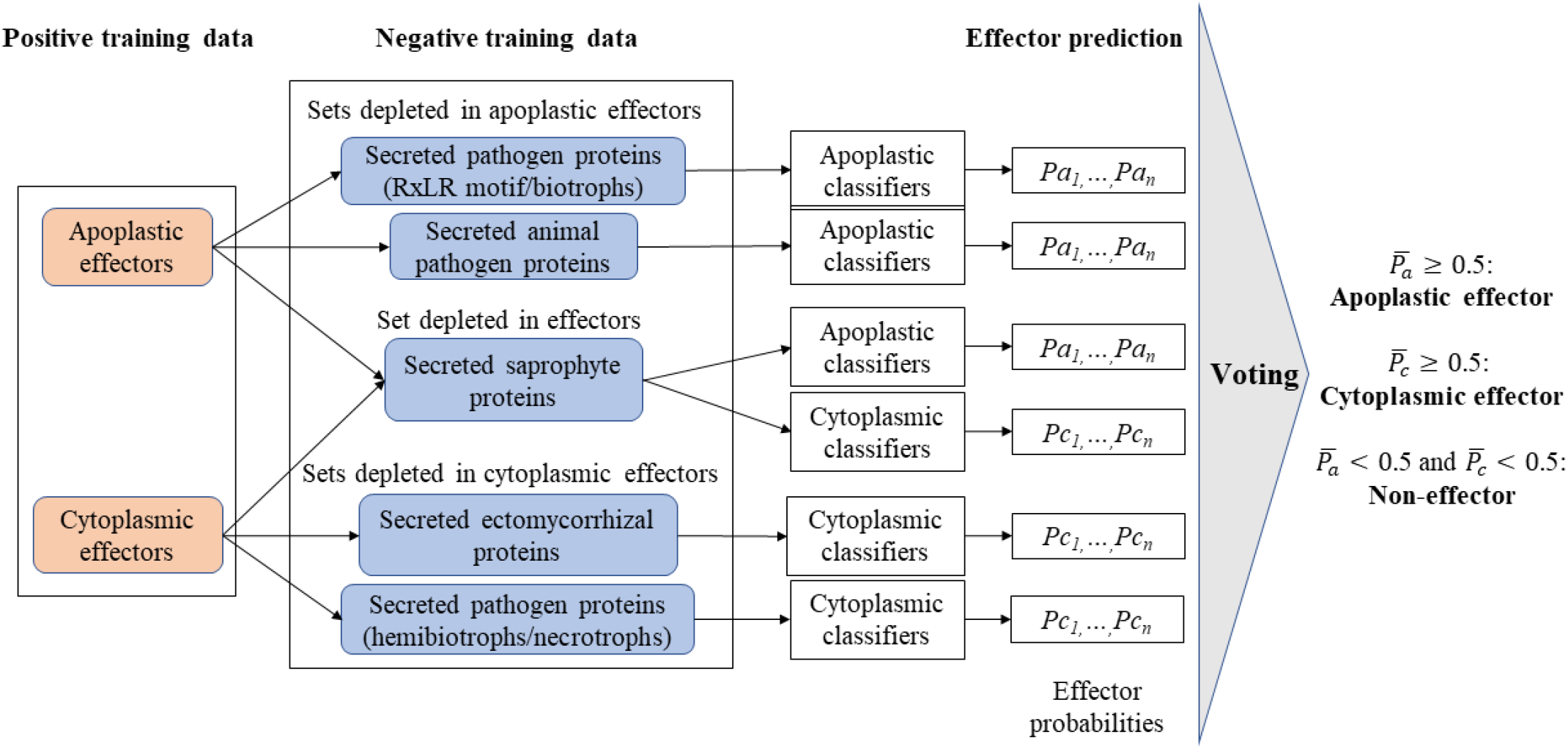
Workflow of EffectorP 3.0. Positive training data are apoplastic and cytoplasmic effectors and negative training data are five different sets of secreted proteins. We trained an ensemble of apoplastic and cytoplasmic classifiers and soft voting is used to arrive at a final prediction for an unseen secreted protein. If a secreted protein is predicted as both apoplastic and cytoplasmic, it is reported as dual-localized and the class with the higher probability is assigned as its most likely localization.

### EffectorP 3.0 has improved accuracy and accurately distinguishes between apoplastic and cytoplasmic localization

To ensure that EffectorP 3.0 is not overfitted and simply memorizes the training data, we collected independent positive and negative test sets. First, a validation set of 48 unseen fungal and oomycete effectors (Supplementary Table S3) was used. EffectorP 3.0 correctly predicts 44 (91.7%) of these as effectors (Table 2). The deep learning tool deepredeff (Kristianingsih & MacLean, 2021) predicts only 56.3% effectors in the validation set correctly. Of the 32 fungal effectors in the set, EffectorP 3.0 and EffectorP-fungi 3.0 both predict 90.6% correctly, whereas EffectorP 2.0 only predicts 74.2% correctly. For the 16 oomycete effectors in the set, EffectorP 3.0 predicts 15 (93.8%) correctly. The EffectorP-fungi 3.0 model only predicts 81.2% of the oomycete effectors correctly, highlighting that a fungal-specific model does not accurately predict oomycete effectors. The same holds for EffectorP 2.0, which was not trained on oomycete effectors and only predicts 43.8% of oomycete effectors correctly. The oomycete effector prediction tool EffectorO (Nur *et al.*, 2021) also predicts a high proportion of the oomycete validation set correctly (14 of 16 effectors, 87.5%). Taken together, EffectorP 3.0 is a more sensitive tool for effector prediction in both fungi and oomycetes than EffectorP 2.0 or deepredeff. We also used a set of secreted fungal saprophyte proteins and secreted proteins from brown algae (*n* = 28,080) as negative test data. This set is depleted for effectors, however it will contain some effector-like proteins. Thus, the false positive rate on this set is a conservative estimation because effector-like proteins are expected to be predicted as effectors by all the prediction methods. EffectorP 3.0 predicts 21.8% as effectors (9.4% cytoplasmic, 12.4% apoplastic) and the balanced accuracy ((sensitivity + specificity)/2) on the positive and negative test sets is ~85%. All other tested methods display lower balanced accuracy (EffectorP 2.0: 81.6%; deepredeff-fungi: 62%; deepredeff-oomycetes: 56.1%; EffectorO: 74.5%).

**Table 2:**
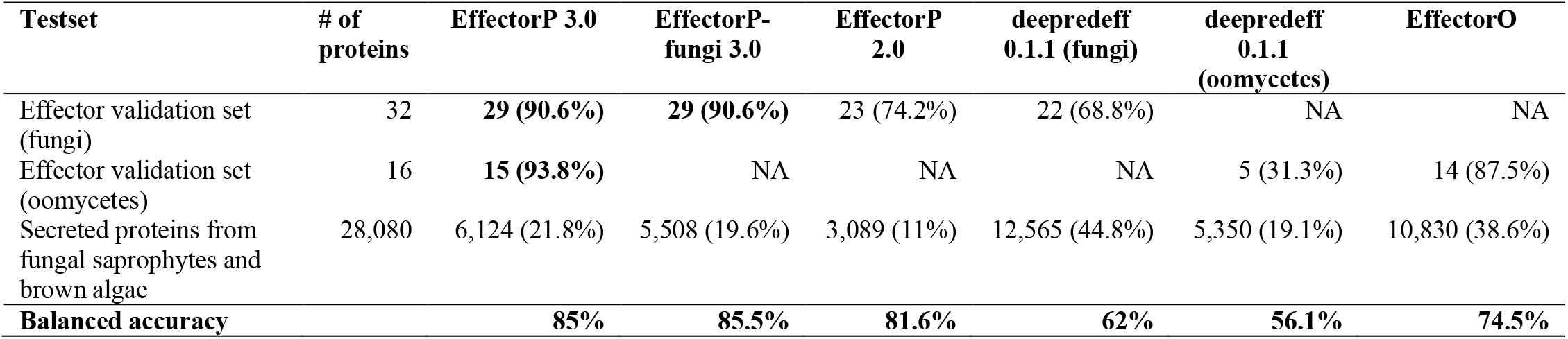
Predicted effectors for different independent test sets. EffectorP 3.0 is the most sensitive method and predicts over 90% of effectors correctly. Balanced accuracy is reported as (sensitivity + specificity)/2. NA: not applicable.

We also estimated EffectorP’s false positive rate for apoplastic and cytoplasmic effector predictions in more detail (Table 3). We first used apoplastic plant proteins, proteins detected in apoplastic proteomics experiments with a predicted signal peptide as well as fungal carbohydrate-active enzymes and secreted saprophyte proteins (*n* = 26,258). These sets are expected to be depleted in cytoplasmic proteins and enriched for secreted apoplastic proteins. From these sets, we estimate that EffectorP 3.0 has a false positive rate for cytoplasmic effector prediction of 8.3% (EffectorP-fungi 3.0: 6.6%). We then collected cytoplasmic plant proteins, human proteins, bacterial type-III effectors as well as fungal non-secreted proteins with a predicted signal peptide, such as those retained in the endoplasmic reticulum (ER) or Golgi apparatus, those directed to the vacuole and those with glycosylphosphatidylinositol (GPI) anchors (*n* = 24,705). All these sets are expected to be depleted in apoplastic effectors. From these sets we estimate that EffectorP 3.0 has a false positive rate for apoplastic effector prediction of 2.5% (EffectorP-fungi 3.0: 2.4%).

**Table 3:**
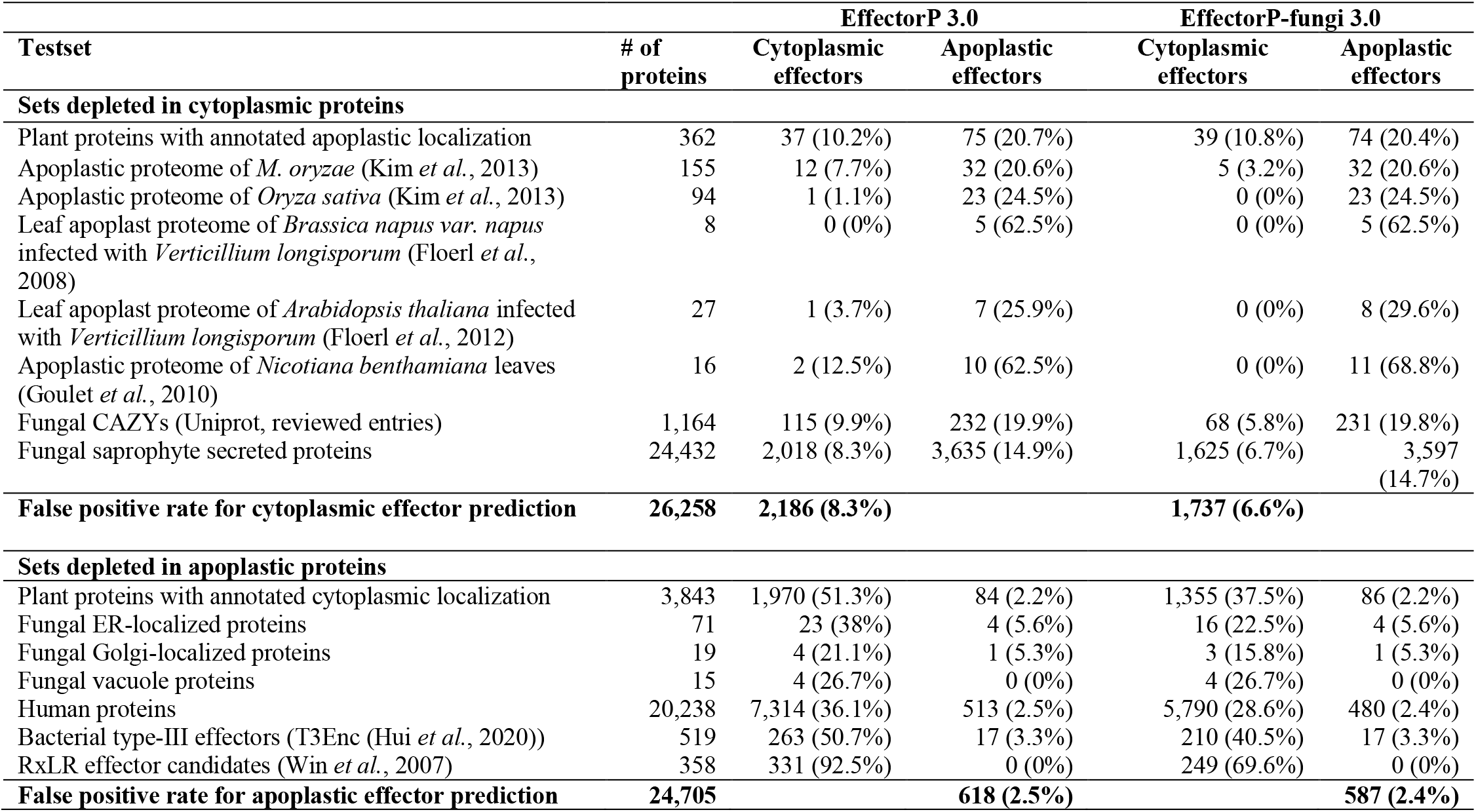
Test sets for assessing the false positive prediction rates for cytoplasmic and apoplastic effector prediction by EffectorP 3.0. In the test sets that are expected to be depleted in cytoplasmic proteins, EffectorP 3.0 has a false positive rate for cytoplasmic effector prediction of 8.3%. A lower false positive rate of 2.5% is achieved by EffectorP 3.0 for apoplastic effector prediction on sets that are depleted in apoplastic proteins.

| Testset | # of proteins | EffectorP 3.0 |  | EffectorP-fungi 3.0 |  |
| --- | --- | --- | --- | --- | --- |
|  |  | Cytoplasmic effectors | Apoplastic effectors | Cytoplasmic effectors | Apoplastic effectors |
| Sets depleted in cytoplasmic proteins |  |  |  |  |  |
| Plant proteins with annotated apoplastic localization | 362 | 37 (10.2%) | 75 (20.7%) | 39 (10.8%) | 74 (20.4%) |
| Apoplastic proteome of *M. oryzae* (Kim *et al.*, 2013) | 155 | 12 (7.7%) | 32 (20.6%) | 5 (3.2%) | 32 (20.6%) |
| Apoplastic proteome of *Oryza sativa* (Kim *et al.*, 2013) | 94 | 1 (1.1%) | 23 (24.5%) | 0 (0%) | 23 (24.5%) |
| Leaf apoplast proteome of *Brassica napus* var. *napus* infected with *Verticillium longisporum* (Floerl *et al.*, 2008) | 8 | 0 (0%) | 5 (62.5%) | 0 (0%) | 5 (62.5%) |
| Leaf apoplast proteome of *Arabidopsis thaliana* infected with *Verticillium longisporum* (Floerl *et al.*, 2012) | 27 | 1 (3.7%) | 7 (25.9%) | 0 (0%) | 8 (29.6%) |
| Apoplastic proteome of *Nicotiana benthamiana* leaves (Goulet *et al.*, 2010) | 16 | 2 (12.5%) | 10 (62.5%) | 0 (0%) | 11 (68.8%) |
| Fungal CAZYs (Uniprot, reviewed entries) | 1,164 | 115 (9.9%) | 232 (19.9%) | 68 (5.8%) | 231 (19.8%) |
| Fungal saprophyte secreted proteins | 24,432 | 2,018 (8.3%) | 3,635 (14.9%) | 1,625 (6.7%) | 3,597 (14.7%) |
| False positive rate for cytoplasmic effector prediction | 26,258 | 2,186 (8.3%) |  | 1,737 (6.6%) |  |
| Sets depleted in apoplastic proteins |  |  |  |  |  |
| Plant proteins with annotated cytoplasmic localization | 3,843 | 1,970 (51.3%) | 84 (2.2%) | 1,355 (37.5%) | 86 (2.2%) |
| Fungal ER-localized proteins | 71 | 23 (38%) | 4 (5.6%) | 16 (22.5%) | 4 (5.6%) |
| Fungal Golgi-localized proteins | 19 | 4 (21.1%) | 1 (5.3%) | 3 (15.8%) | 1 (5.3%) |
| Fungal vacuole proteins | 15 | 4 (26.7%) | 0 (0%) | 4 (26.7%) | 0 (0%) |
| Human proteins | 20,238 | 7,314 (36.1%) | 513 (2.5%) | 5,790 (28.6%) | 480 (2.4%) |
| Bacterial type-III effectors (T3Enc (Hui *et al.*, 2020)) | 519 | 263 (50.7%) | 17 (3.3%) | 210 (40.5%) | 17 (3.3%) |
| RxLR effector candidates (Win *et al.*, 2007) | 358 | 331 (92.5%) | 0 (0%) | 249 (69.6%) | 0 (0%) |
| False positive rate for apoplastic effector prediction | 24,705 | 618 (2.5%) |  | 587 (2.4%) |  |

Finally, we applied EffectorP 3.0 to the cytoplasmic and apoplastic effectors that have experimental evidence for their localization from both the training and validation sets (Table 4). 80% of the 110 cytoplasmic effectors are predicted correctly as cytoplasmic with EffectorP 3.0. The eight cytoplasmic effectors incorrectly predicted as apoplastic are RTP1 (*Uromyces viciae-fabae*), PpEC23 (*Phakopsora pachyrhizi*), Foa3 (*Fusarium oxysporum* Fo5176), AvrPiz-t (*Magnaporthe oryzae*), SsSSVP1 (*Sclerotinia sclerotiorum*), HvEC_016 (*Hemileia vastatrix*), AvrPm2 (*Blumeria graminis* f. sp. *tritici*) and AVRa9 (*Blumeria graminis* f. sp. *hordei*). These effectors range in sequence length from 102-291 aas with a cysteine number of 3-20 (cysteine content: 2.3%-6.9%). Thus, a high cysteine content together with a small size could be the reason for the apoplastic mis-predictions. A further seven cytoplasmic effectors are predicted as dual-localized effectors, with apoplastic as the class with highest probability (AvrP123 (*Melampsora lini*), SIX5 (*Fusarium oxysporum* f. sp. *lycopersici*), AvrLm4-7 (*Leptosphaeria maculans*), AVR-Pii and AvrPi9 (*Magnaporthe oryzae*), ToxA (*Parastagonospora nodorum*), AvrSr27 (*Puccinia graminis* f. sp. *tritici*)). These effectors range in sequence length from 70-143 aas with a cysteine number of 2-15 (cysteine content: 1%-10.4%). Whilst these effectors are incorrectly predicted with apoplastic as their most likely localization, they are still being predicted as cytoplasmic albeit with a lower probability. For example, AvrP123 is a cytoplasmic effector from the rust fungus *Melampsora lini* with 11 cysteines and a small size of 117 aas (9.4% cysteine content) and the cysteine residues are involved in zinc binding instead of disulfide bond formation (Zhang *et al.*, 2018). EffectorP 3.0 predicts AvrP123 as cytoplasmic with probability 0.77 and apoplastic with probability 0.8, thus still recognizing the cytoplasmic signal despite the high cysteine content. Similarly, AvrSr27 is a cytoplasmic effector from the rust fungus *Puccinia graminis* f.sp. *tritici* with 15 cysteines and a small size of 144 aas (10.4% cysteine content) (Upadhyaya *et al.*, 2021). Again, EffectorP 3.0 recognizes the cytoplasmic signal and predicts AvrSr27 as cytoplasmic with probability 0.72 and apoplastic with probability 0.78. Finally, we note that EffectorP 3.0 predicts 51.3% of cytoplasmic plant proteins as cytoplasmic effectors. This suggests that EffectorP 3.0 will recognize a cytoplasmic signal also in intracellular proteins. Thus, for accurate effector prediction EffectorP should only be applied to proteins with computational or experimental evidence for secretion.

**Table 4:** Predicted localizations for cytoplasmic and apoplastic effectors that have experimental evidence for their localization from both the training and validation sets. EffectorP 3.0 predicts ~80% of apoplastic and cytoplasmic effector localizations correctly, whereas ApoplastP 1.0 only predicts the localization of ~77% of the apoplastic effectors correctly.

|  |  | EffectorP 3.0 |  | EffectorP-fungi 3.0 |  | ApoplastP 1.0 |
| --- | --- | --- | --- | --- | --- | --- |
|  | # of proteins | Cytoplasmic effectors | Apoplastic effectors | Cytoplasmic effectors | Apoplastic effectors | Apoplastic proteins |
| **Apoplastic effectors with experimental support from validation and training sets** | 57 | 7 (12.3%) | **46 (80.7%)** | 12 (10.9%) | 44 (77.2%) | 44 (77.2%) |
| **Cytoplasmic effectors with experimental support from validation and training sets** | 110 | **88 (80%)** | 15 (13.6%) | 77 (70%) | 9 (15.8%) | 16 (14.5%) |

### Cytoplasmic effectors are enriched for positively charged residues

We further investigated which protein features might be most important for apoplastic and cytoplasmic effector predictions in fungi and oomycetes. For this, we selected the most discriminative features that separate effectors from secreted proteins in saprophytes and brown algae. For fungal cytoplasmic effectors, the four most discriminative features are an enrichment for the positively charged amino acid lysine, low molecular weight, enrichment for amino acids with high flexibility and enrichment for positively charged residues. Similarly, for oomycete cytoplasmic effectors, the four most discriminative features are an enrichment in the positively charged amino acid lysine, an enrichment for positively charged residues, depletion in surface exposed residues and an enrichment for residues that frequently occur in alpha helices. An enrichment for positively charged amino acids such as lysine appears to be a unifying feature for both fungal and oomycete cytoplasmic effectors, but not for apoplastic effectors (Figure 2). We further investigated the lysine content in the signal peptide regions (first 20 aas), N-terminal regions (the first third of the sequence after the signal peptide region) and the C-terminal regions (last two-thirds of the sequence). Both fungal and oomycete cytoplasmic effectors are enriched for lysines in the C-terminal region, suggesting that this is a feature of the mature protein after cleavage of the signal peptide, and after the RxLR region in oomycete effectors (Figure 3). This feature could be related to effector function in the cytoplasm or the requirements for translocation.

**Figure 2:**
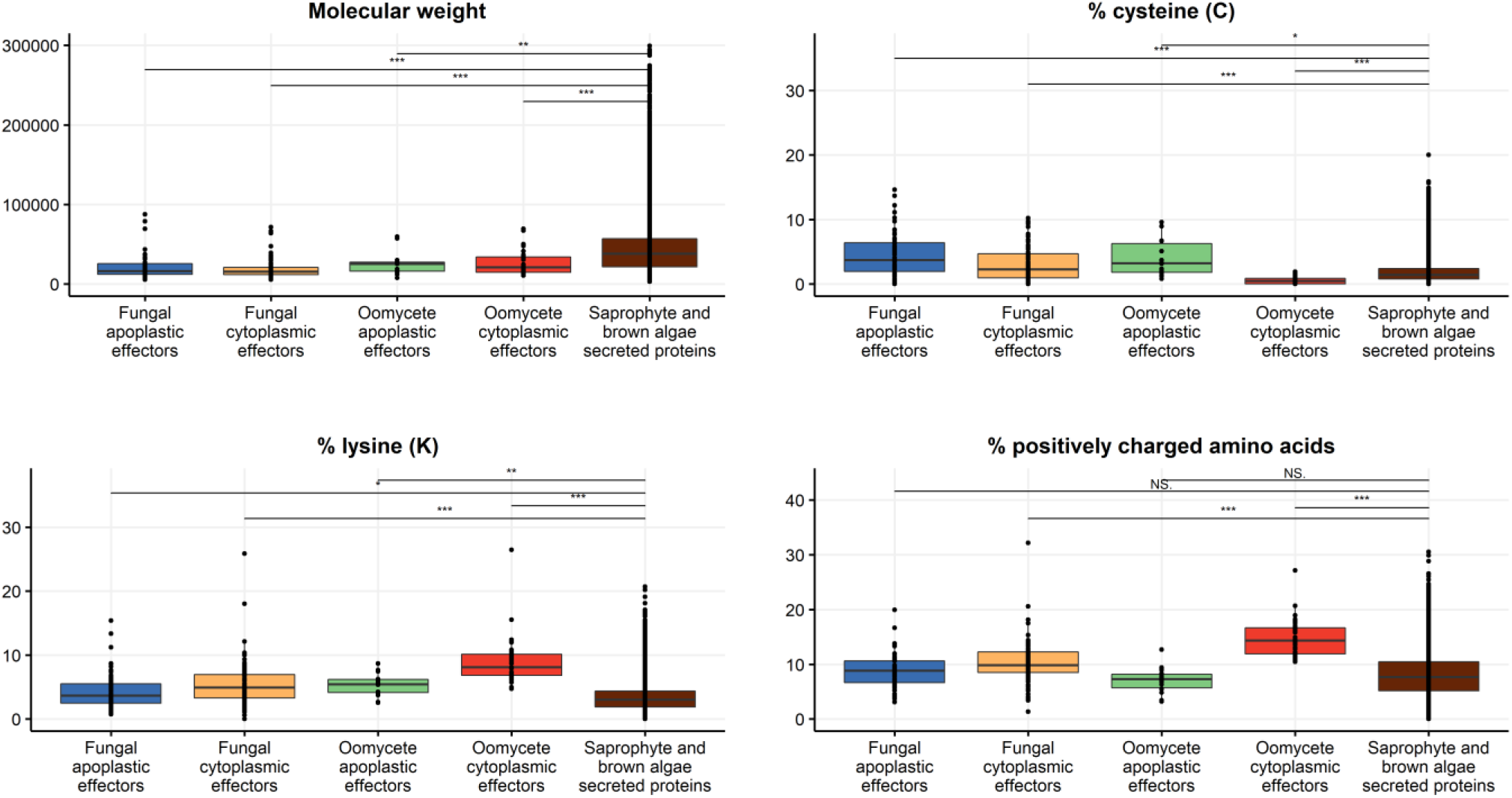
The most discriminate features in fungal and oomycete effector prediction. Fungal and oomycete effectors have on average lower molecular weight than saprophyte/brown algae secreted proteins. A unifying feature for fungal and oomycete cytoplasmic effectors is an enrichment for positively charged amino acids, particularly lysines.

**Figure 3:**
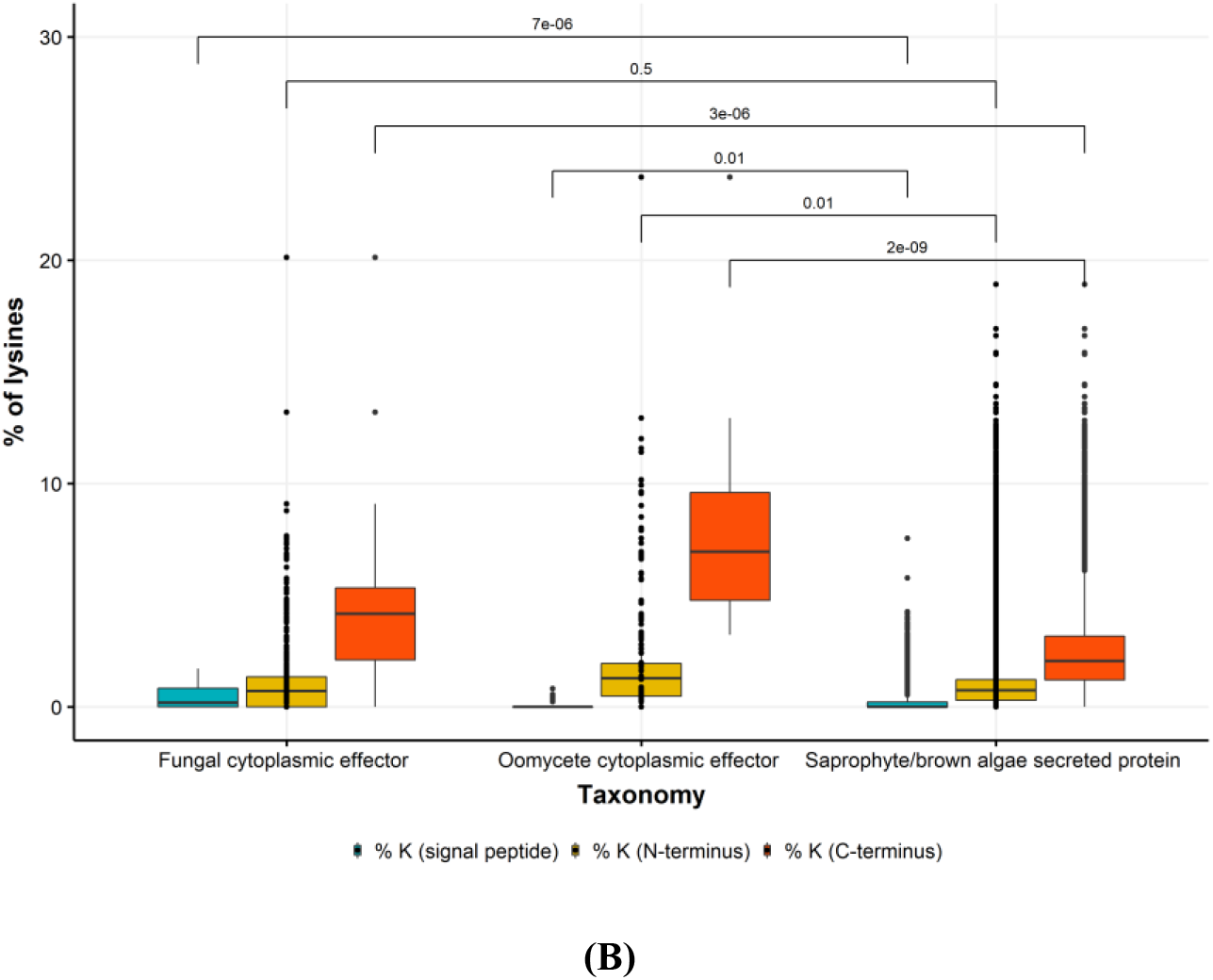
The C-terminal regions of fungal and oomycete effectors (last two-thirds of the sequence) are enriched for lysine residues.

Both apoplastic and cytoplasmic effectors have on average lower molecular weight than secreted non-pathogen proteins. Apoplastic fungal and oomycete effectors are both enriched for cysteines, which was not observed for oomycete cytoplasmic effectors (Figure 2). An enrichment for cysteines is also observed for fungal cytoplasmic effectors. Taken together, this suggests that low molecular weight and a high cysteine content is generally a unifying feature for apoplastic effectors and low molecular weight and a high lysine content is a unifying feature for cytoplasmic effectors. Thus, selection of fungal cytoplasmic effector candidates for experimental validation should not be based on low molecular weight and a high cysteine content alone.

### Sets of infection‐induced proteins are enriched for predicted effectors

Effectors are expressed during infection and thus, secreted proteins encoded by genes that are expressed during infection are expected to be enriched for predicted effectors. We used several expression data sets from the literature to assess if EffectorP 3.0 leads to improved effector predictions. All infection-induced effector candidate sets are enriched for predicted effectors by EffectorP 3.0 when compared to effector predictions on whole secretomes (Table 5). Furthermore, the predicted apoplastic and cytoplasmic localization of effector candidates correspond well with the corresponding infection stage. For example, genes encoding for secreted proteins that are up-regulated in biotrophic invasive hyphae in *Magnaporthe oryzae* (Mosquera et al., 2009) are enriched for predicted cytoplasmic effectors, while *in planta* induced small secreted apoplastic effector candidates in *Cladosporium fulvum* (Mesarich *et al*., 2017) are enriched for predicted apoplastic effectors. We then used gene expression data of the obligate biotrophic fungal pathogen *Puccinia graminis* f. sp. *tritici* 21-0 to investigate expression of predicted apoplastic and cytoplasmic effectors (Chen *et al.*, 2017). This time course covers wheat infection from 2-7 days post infection (dpi) as well as *in vitro* germinated spores and haustorial tissue (Upadhyaya *et al.*, 2015). Rust haustoria enable nutrient uptake as well as the delivery of effectors into the host plant cell and are thus expected to be enriched for cytoplasmic effectors. In contrast, spores germinate on the leaf surface at the start of the infection cycle and are thus expected to be depleted for cytoplasmic effectors. We collected secreted proteins that are up-regulated in germinated spores compared to haustorial tissue (*n* = 841) and secreted proteins that are up-regulated in haustorial tissue compared to germinated spores (*n* = 1,732). EffectorP 3.0 predicts 22.1% and 21.9% of the secreted proteins up-regulated in germinated spores as cytoplasmic and apoplastic effectors, respectively. In contrast, EffectorP 3.0 predicts 55.7% and 18.5% of the secreted proteins up-regulated in haustoria as cytoplasmic and apoplastic effectors, respectively. Thus, as expected secreted proteins up-regulated in rust haustorial tissue are enriched for predicted cytoplasmic effectors with EffectorP 3.0.

**Table 5:** Infection-induced genes encoding secreted proteins are enriched for predicted effectors by EffectorP 3.0.

| Testset | # of proteins | EffectorP 3.0 |  |
| --- | --- | --- | --- |
|  |  | % predicted effectors (apoplastic/cytoplasmic) | % predicted effectors in secretome (apoplastic/cytoplasmic) |
| *Colletotrichum higginsianum*: biotrophy-associated effector candidates (Kleemann et al., 2012) | 102 | 62.7% (48.4%/51.6%) | 26.6% (67.3%/32.7%) |
| *Cladosporium fulvum*: *in planta* induced small secreted apoplastic effector candidates (Mesarich et al., 2017) | 75 | 94.7% (85.9%/14.1%) | 29.5% (61.4%/38.6%) |
| *Magnaporthe oryzae*: genes with $\geq 50$-fold differential expression in biotrophic invasive hyphae (Mosquera et al., 2009) | 15 | 100% (33.3%/66.7%) | 38.9% (49.4%/50.6%) |
| *Blumeria graminis* f. sp. *hordei*: Candidates for Secreted Effector Proteins (CSEPs) (Pedersen et al., 2012) | 491 | 81.5% (18%/82%) | 48.7% (24.3%/75.7%) |
| *Laccaria bicolor*: ectomycorrhiza-regulated small secreted proteins (MiSSPs) (Martin et al., 2008) | 21 | 85.7% (50%/50%) | 22.5% (55.8%/44.2%) |
| *Zymoseptoria tritici* candidate effectors (Kettles et al., 2017) | 63 | 76.2% (68.8%/31.2%) | 32.2% (57.3%/42.7%) |
| *Ustilago maydis* effector candidates (Tollot et al., 2016) | 198 | 49% (30.9%/69.1%) | 35.7% (37.7%/62.3%) |
| *Leptosphaeria maculans* highly expressed early effector candidates (Gervais et al., 2017) | 49 | 83.7% (41.5%/58.5%) | 24.3% (58.9%/41.1%) |
| *Leptosphaeria maculans* highly expressed late effector candidates (Gervais et al., 2017) | 50 | 62% (48.4%/51.6%) | 24.3% (58.9%/41.1%) |
| *Bremia lactucae* cell death inducing effector candidates (Wood et al., 2020) | 11 | 72.7% (0%/100%) | 51.6% (20.2%/79.8%) |

### Obligate biotroph and animal parasite secretomes are highly enriched for predicted cytoplasmic effectors

We predicted apoplastic and cytoplasmic effectors in 92 fungal and oomycete secretomes (Table 6, Supplementary Data S1). The highest proportions of cytoplasmic effectors are predicted in the secretomes of the animal parasites *Encephalitozoon cuniculi* (75.4%) and *Enterocytozoon bieneusi* (72%), the animal pathogen *Batrachochytrium dendrobatidis* (55.6%), the arbuscular mycorrhizal fungus *Rhizophagus clarus* (52%) and the obligate biotrophic pathogens *Albugo laibachii* (49.4%) and *Puccinia graminis* f. sp. *tritici* (44.7%). The highest proportions of apoplastic effectors are predicted in the secretomes of the hemibiotrophic pathogen *Venturia inaequalis* (24%), the necrotrophic pathogen *Sclerotinia sclerotiorum* (23.7%), the arbuscular mycorrhizal fungus *Gigaspora rosea* (22.5%), the obligate biotrophic pathogens *Puccinia striiformis* f. sp. *tritici* (21.3%) and *P. graminis* f. sp. *tritici* (20.5%) as well as the hemibiotrophic pathogen *Fusarium graminearum* (19.8%). Overall, the secretomes of obligate biotrophs, biotrophic pathogens, hemibiotrophic pathogens, arbuscular mycorrhizal fungi and animal pathogens/parasites have higher proportions of predicted cytoplasmic effectors than apoplastic effectors in their secretomes (Table 6). As expected, necrotrophic pathogens and ectomycorrhizal fungi are enriched for predicted apoplastic effectors.

**Table 6:** Predicted effectors in fungal and oomycete secretomes. Apoplastic and cytoplasmic effectors were predicted with EffectorP 3.0 and with EffectorP-fungi 3.0. Significance between cytoplasmic effector predictions with EffectorP 3.0 and EffectorP-fungi 3.0 were tested with paired t-tests (*, < 0.05; **, < 0.01; ***, < 0.001; ^NS^, not significant). Significantly higher numbers of cytoplasmic effectors were predicted with the EffectorP 3.0 model trained on both fungi and oomycete cytoplasmic effectors in all groups except animal parasites.

| Classification | # of genomes | Average # of secreted proteins | EffectorP 3.0 predictions |  | EffectorP-fungi 3.0 predictions |  |
| --- | --- | --- | --- | --- | --- | --- |
|  |  |  | % cytoplasmic effectors | % apoplastic effectors | % cytoplasmic effectors | % apoplastic effectors |
| Obligate biotrophs | 15 | 1,770 | **38.7%\*** | 12.1% | 32.5% | 11.3% |
| Biotrophic pathogens | 2 | 438 | **23%\*** | 11.3% | 18.4% | 11.2% |
| Hemibiotrophic pathogens | 16 | 1,474 | **18.2%\*\*** | 17.3% | 15.1% | 16.7% |
| Necrotrophic pathogens | 9 | 1,101 | 12.4%*** | **17.9%** | 9.7% | 17.6% |
| Ectomycorrhizal fungi | 11 | 1,201 | 9.7%*** | **12.9%** | 8.3% | 12.7% |
| Arbuscular mycorrhizal fungi | 5 | 861 | **40.6%\*** | 16% | 35.4% | 15.2% |
| Saprophytic fungi | 21 | 1,090 | 8.3%*** | **14.4%** | 6.9% | 14.2% |
| Yeasts | 5 | 372 | **15.6%\*\*** | 9.4% | 11.6% | 9.1% |
| Brown algae | 4 | 1,254 | **13.4%\*\*** | 4.1% | 8% | 3.9% |
| Animal fungal pathogens | 7 | 558 | **23.4%\*** | 8.3% | 19.2% | 8.2% |
| Animal parasite | 2 | 106 | 73.7%<sup>NS</sup> | 2.7% | 63.8% | 3.2% |

Non-pathogenic fungi that colonize plants include endophytes as well as ectomycorrhizal and arbuscular mycorrhizal fungi, which establish close symbiotic association with plants. Symbiotic fungi such as arbuscular mycorrhizal fungi have dedicated feeding structures (arbuscules) and also secrete cytoplasmic effector proteins to suppress plant defense and promote symbiosis (Kloppholz *et al.*, 2011; Plett *et al.*, 2011). Indeed, EffectorP 3.0 predicts a high average proportion of cytoplasmic effectors (40.6%) for the five tested arbuscular mycorrhizal fungal secretomes. In contrast, only 8.3% predicted cytoplasmic effectors occur on average in the saprophyte secretomes. However, we found a substantially higher proportion of predicted cytoplasmic effectors in yeasts (15.6%). We further investigated the high-quality functional annotations of the predicted effectors of the yeast *Saccharomyces cerevisiae* (Supplementary Tables S5 and S6). Of the 268 predicted secreted proteins, EffectorP 3.0 predicts 46 cytoplasmic effectors (17.2%) and 29 apoplastic effectors (10.8%). Most of the predicted cytoplasmic effectors have annotated intracellular subcellular localizations, suggesting that EffectorP 3.0 is predicting an intracellular signal and these proteins are non-secreted false positives from SignalP 4.0 (Petersen *et al.*, 2011). This underlines that accurate secretion prediction is a pre-requisite for accurate EffectorP predictions. Current versions of SignalP (5.0, 6.0) might reduce the number of false positive secreted proteins, however this might come at the cost of missing true positive effectors in the secretomes (Sperschneider *et al.*, 2015b).

## Discussion

Plant diseases and pests are a major threat to food security. Plant pathogens cause disease through secreted effector proteins and their identification from genomic sequences is crucial for enhancing host-plant resistance and for improving crop resistance (Vleeshouwers & Oliver, 2014). With the ever-increasing number of high-quality genome sequences, accurate computational prediction of effectors for subsequent experimental validation has become the bottleneck. Existing effector prediction methods return hundreds of candidates and can suffer from high false positive rates (Sperschneider *et al.*, 2015), leading to low experimental validation success rates and a limited understanding of effector biology.

Fungal effector proteins are diverse in sequence which complicates their prediction. Early approaches to fungal effector prediction were reliant on filtering secretomes for proteins of small size and high cysteine content. Such user-driven selection is generally of low accuracy and has been detrimental for advancing our understanding of effector biology as it is unable to uncover exceptions to the rules. A contrasting data-driven approach is machine learning that can automatically identify relevant patterns in data. A machine learning model can be trained to recognize a particular concept based on its features in observed data and can then be applied to identify new instances of the concept in unseen data. EffectorP is such a machine learning prediction tool for fungal effector prediction (Sperschneider *et al.*, 2016, 2018). However, current effector prediction methods such as EffectorP 2.0 (Sperschneider *et al.*, 2018a), deepredeff (Kristianingsih & MacLean, 2021) or EffectorO (Nur *et al.*, 2021) give a yes-or-no answer as to whether a protein is a likely effector and do not deliver vital information for prioritization: Is the effector extracellular and functions in the plant apoplast or is it cytoplasmic and enters plant cells? In oomycetes, the presence of conserved sequence motifs such as the RxLR motif have been used extensively for cytoplasmic effector prediction. However, the biological function of the RxLR motif in effector translocation has been debated (Ellis & Dodds, 2011) (Wawra *et al.*, 2017). Furthermore, the RxLR motif is not conserved outside *Phytophthora* species and downy mildews and can thus not be used for effector prediction alone in other oomycete species (Wood *et al.*, 2020) (Ai *et al.*, 2020). Whilst in oomycetes sequence motif and Hidden Markov Model searches are well-established methods to predict cytoplasmic effectors, this will miss *bona fide* effectors that do not carry such signatures. Furthermore, apoplastic effector prediction in oomycetes is an unresolved problem.

Here, we take advantage of differences in apoplastic and cytoplasmic environments that allow for accurate prediction of pathogen proteins localizing to these compartments. EffectorP 3.0 can predict if a secreted protein is an apoplastic effector, a cytoplasmic effector or a non-effector. On our evaluation sets, we found that EffectorP 3.0 has higher accuracy than EffectorP 2.0, deepredeff or EffectorO. It is also more accurate in predicting apoplastic localization in effectors than ApoplastP (Sperschneider *et al.*, 2018b). We found that cytoplasmic effectors are enriched for positively charged amino acids in the C-terminus, in both fungi and oomycetes. In contrast, the deep learning tool deepredeff found the most discriminative signal for effector prediction in the first 25 aas of the proteins, i.e. in the signal peptide region. In our tests, deepredeff has a high false positive rate in non-pathogen secretomes, suggesting that the signal peptide region alone is unlikely the discriminative signal for effector function. A high cysteine content was an important component of prediction for apoplastic effectors. However, in some cases this signal led to cysteine-rich cytoplasmic effectors being assigned a high probability of apoplastic localization. Some cytoplasmic effectors, such as AvrP123 (Zhang *et al.*, 2018) contain high numbers of cysteine residues that can be involved in metal ion coordination, rather than disulfide bond formation. This potential for mis-prediction of localization should be kept in mind when interpreting EffectorP 3.0 results for cysteine-rich effector candidates. In the future, larger training data sets and additional experimental evidence for effector localization will allow for even more accurate predictions, as well as for investigations if any protein features are involved in effector translocation.

It is important to apply EffectorP 3.0 to secreted proteins only, and not to non-secreted proteins. EffectorP 3.0 will predict a substantial proportion of intracellular, non-secreted proteins as cytoplasmic effectors. For example, over half of plant cytoplasmic proteins are predicted as cytoplasmic effectors. This is not unexpected, as cytoplasmic effectors localize to the same environment as plant intracellular proteins. Again, accurate secretion prediction is a pre-requisite for EffectorP 3.0. Lastly, whilst EffectorP 3.0 returns probabilities for the likelihood that a secreted protein is an effector, we recommend that these are not over-interpreted. Ranking effector candidates according to their expression during infection with RNAseq data, or through structural similarity to known effectors are recommended practices for effector candidate prioritization.

## Materials and Methods

### Curation of positive and negative training data

Effector sequences were collected through literature searches. For the negative training sets, we predicted secretomes from a wide range of fungal and oomycete genomes (Supplementary Table S1) with SignalP 4.1 (-t euk -u 0.34 -U 0.34) (Petersen *et al.*, 2011) and TMHMM 2.0 (Krogh *et al.*, 2001). A fungal protein was called secreted if it was predicted to have a signal peptide and has no transmembrane domains. A secreted oomycete protein was defined as an RxLR protein if it contains the RxLR motif in the first 100 amino acids. We curated five secreted protein sets 1) a homology-reduced set of secreted proteins with an RxLR motif from plant-pathogenic oomycetes together with secreted proteins from biotrophic or hemibiotrophic fungi; 2) a homology-reduced set of secreted proteins without an RxLR motif from plant-pathogenic oomycetes together with secreted proteins from necrotrophic or hemibiotrophic fungi; 3) a set of secreted fungal saprophyte proteins and secreted proteins from brown algae; 4) a set of secreted fungal ectomycorrhizal proteins; and 5) a set of secreted proteins from animal-pathogenic fungi and animal-pathogenic oomycetes. Proteins in the negative sets that share sequence homology to an effector in the positive training set were excluded and protein sets were homology-reduced, both with cd-hit 4.8.1 (-c 0.9) (Fu *et al.*, 2012).

### Machine learning model training and feature extraction

We used a variety of protein features in classification: 1) amino acid frequencies; 2) molecular weight; 3) % of positively charged amino acids (K, R) in the sequence; 4) % of negatively charged amino acids (D, E) in the sequence; 5) average surface exposure of amino acids in the sequence (Janin, 1979); 6) average hydrophobicity of amino acids in the sequence (Fauchere & Pliska, 1983); 7) average polarity of amino acids in the sequence (Zimmerman *et al.*, 1968); 8) average flexibility of amino acids in the sequence (Vihinen *et al.*, 1994); 9) frequency of aromatic amino acids (F, H, W, Y) in the sequence; 10) frequency of polar amino acids (D, E, H, K, N, Q, R, S, T, Z) in the sequence; 11) average disorder propensity of amino acids in the sequence (Dunker *et al.*, 2001); 12) average bulkiness of amino acids in the sequence (Zimmerman *et al.*, 1968); 13) average alpha helix frequency for amino acids in the sequence (Nagano, 1973); 14) average beta sheet frequency for amino acids in the sequence (Nagano, 1973); and 15) average coil frequency for amino acids in the sequence (Nagano, 1973). The WEKA tool box (version 3.8.4) was used to train and evaluate the performance of different machine learning classifiers (Hall *et al*., 2000).

For the effector prediction models, we took randomly selected samples of each negative training data set to give a ratio of 3 : 1 to the number of positive training examples. We then used WEKA to train Naïve Bayes classifiers on each of the negative datasets with the same positive training set. We then repeated this procedure and trained C4.5 decision trees (J48 model in WEKA) on each of the negative datasets. For each set of 100 models, we selected the best‐performing models as those with the highest negative predictive value (NPV: True negatives/(True negatives + False negatives)). Five Naïve Bayes classifiers and five C4.5 decision trees that discriminate between apoplastic effectors and secreted pathogen proteins; five Naïve Bayes classifiers and five C4.5 decision trees that discriminate between apoplastic effectors and secreted saprophyte/brown algae proteins; five Naïve Bayes classifiers and five C4.5 decision trees that discriminate between apoplastic effectors and secreted animal-pathogenic pathogen proteins; five Naïve Bayes classifiers and five C4.5 decision trees that discriminate between cytoplasmic effectors and secreted pathogen proteins; five Naïve Bayes classifiers and five C4.5 decision trees that discriminate between cytoplasmic effectors and secreted saprophyte/brown algae proteins; five Naïve Bayes classifiers and five C4.5 decision trees that discriminate between cytoplasmic effectors and secreted fungal ectomycorrhizal proteins. The ensemble classifier called EffectorP 3.0 returns a final prediction using a soft voting approach, which predicts the class label based on average probabilities for ‘apoplastic effector’, ‘cytoplasmic effector and ‘non‐effector’ calculated by each classifier. Soft voting then returns the class with the highest average probability as the result. A protein is classified as an apoplastic effector if the average probability for the class ‘apoplastic effector’ is higher than the average probability for the class ‘cytoplasmic effector’ and the class ‘non‐effector’. A protein is classified as a cytoplasmic effector if the average probability for the class ‘cytoplasmic effector’ is higher than the average probability for the class ‘apoplastic effector’ and the class ‘non‐effector’. If a secreted protein is predicted as both an apoplastic and cytoplasmic effector, EffectorP 3.0 reports it as dual-localized and the class with the highest probability as its likely localization. As the number of cytoplasmic fungal effectors is sufficiently large, we also trained a fungal-specific cytoplasmic model with the same negative data sets and apoplastic model as above (EffectorP-fungi 3.0). Weka’s CorrelationAttributeEval + Ranker method were used to find the most discriminative features for classification.

All plots were produced using Ggplot2 (Wickham, 2009) and statistical significance was assessed with *t*-tests using the ggsignif package (https://cran.r-project.org/web/packages/ggsignif/index.html). Significance thresholds according to *t*-test are: NS, not significant; *, < 0.05; **, < 0.01; ***, < 0.001.

### Collection of validation sets

Validation sets were collected from UniProt with the following search terms. Plant proteins with annotated apoplastic localization: “locations:(location:“Apoplast [SL-0019]”) length:[50 TO *] taxonomy:“Viridiplantae [33090]” AND reviewed:yes”; Plant proteins with annotated cytoplasmic localization: “taxonomy:“Viridiplantae [33090]” locations:(location:“Cytoplasm [SL-0086]”) length:[50 TO *] AND reviewed:yes”; Fungal CAZYs: “taxonomy:“ Fungi [4751]” database: cazy AND reviewed:yes”. All other sets were collected from the literature and are available at https://github.com/JanaSperschneider/EffectorP-3.0-Data.

### RNAseq differential expression analysis

RNA reads from germinated spores and haustorial tissue of *Pgt* 21-0 were obtained from National Center for Biotechnology Information (NCBI) BioProject PRJNA253722 (Upadhyaya *et al.*, 2015) while reads from an infection timecourse (Chen *et al.*, 2017) were obtained from NCBI BioProject PRJNA415866. Salmon v.1.1.0 (Patro *et al.*, 2017) was used to align reads to the *Pgt* 21-0 transcripts and to estimate transcript abundances in each sample (salmon index –keepDuplicates and salmon quant –validateMappings). We used tximport and DESeq2 (Love *et al.*, 2014) to assess gene differential expression.

## Acknowledgments

JS is supported by an Australian Research Council (ARC) Discovery Early Career Researcher Award (DE190100066).

