## Supplementary Table for "EffectorP 3.0: prediction of apoplastic and cytoplasmic effectors in fungi and oomycetes"

**Supplementary Table S1**: The genomes used for training of EffectorP 3.0.

| Description | Genomes used |
| --- | --- |
| Secretomes from biotrophic plant-pathogenic oomycetes | *Albugo laibachii* (Kemen et al., 2011)  *Bremia lactucae* (Wood et al., 2020)  *Hyaloperonospora arabidopsidis* (Baxter et al., 2010)  *Peronospora effusa* (Fletcher et al., 2018) |
| Secretomes from hemibiotrophic plant-pathogenic oomycetes | *Phytophthora capsici* (Lamour *et al.*, 2012)  *Phytophthora infestans* (Haas *et al.*, 2009)  *Phytophthora parasitica* (Genbank GCA_000247585.2)  *Phytophthora ramorum* (Tyler *et al.*, 2006)  *Phytophthora sojae* (Tyler *et al.*, 2006) |
| Secretomes from necrotrophic plant-pathogenic oomycetes | *Pythium ultimum* (Levesque *et al.*, 2010) |
| Secretomes from brown algae | *Cladosiphon* *okamuranus* (Nishitsuji et al., 2016)  *Ectocarpus siliculosus* (Cock et al., 2010)  *Nemacystus decipiens* (Nishitsuji et al., 2019)  *Undaria pinnatifida* (Shan et al., 2020) |
| Secretomes from animal-pathogenic oomycetes | *Saprolegnia parasitica (*Jiang et al., 2013) |
| Secretomes from biotrophic plant-pathogenic fungi | *Blumeria graminis* f. sp. *hordei* (Muller et al., 2019)  *Blumeria graminis* f. sp*. tritici* (Frantzeskakis et al., 2018)  *Erysiphe necator* (Jones et al., 2014)  *Melampsora lini* (Sperschneider et al., unpublished)  *Puccinia graminis* f.sp*. tritici* (Li et al., 2019)  *Puccinia striiformis* f.sp*. tritici* (Schwessinger et al., 2018)  *Ustilago maydis* (Kämper et al., 2006)  *Ustilago hordei* (Laurie et al., 2012) |
| Secretomes from hemibiotrophic plant-pathogenic fungi | *Cladosporium fulvum* (de Wit et al., 2012)  *Cochliobolus sativus* (Ohm et al., 2012)  *Colletotrichum graminicola* (O'Connell et al., 2012)  *Colletotrichum higginsianum* (Zampounis et al., 2016)  *Fusarium graminearum* (Cuomo et al., 2007)  *Fusarium oxysporum f. sp. lycopersici* (Ma et al., 2010)  *Leptosphaeria maculans* (Rouxel et al., 2011)  *Magnaporthe oryzae* (Dean et al., 2005)  *Venturia inaequalis* (Deng et al., 2017)  *Verticillium dahlia* (Klosterman et al., 2011)  *Zymoseptoria tritici* (Goodwin et al., 2011) |
| Secretomes from necrotrophic plant-pathogenic fungi | *Alternaria alternata* (Zeiner et al., 2016)  *Alternaria brassicicola* (Dang et al., 2015)  *Ascochyta rabiei* (Verma et al., 2016)  *Botrytis cinerea* (Amselem et al., 2011)  *Cochliobolus miyabeanus* (Condon et al., 2013)  *Cochliobolus victoriae* (Condon et al., 2013)  *Sclerotinia sclerotiorum* (Amselem et al., 2011)  *Parastagonospora nodorum* (Hane et al., 2007) |
| Secretomes from ectomycorrhizal fungi | *Amanita muscaria* (Kohler et al., 2015)  *Hebeloma cylindrosporum* (Kohler et al., 2015)  *Laccaria amethystine* (Kohler et al., 2015)  *Laccaria bicolor* (Martin et al., 2008)  *Paxillus involutus* (Kohler et al., 2015)  *Paxillus rubicundulus* (Kohler et al., 2015)  *Piloderma olivaceum* (Kohler et al., 2015)  *Pisolithus microcarpus* (Kohler et al., 2015)  *Pisolithus tinctorius* (Kohler et al., 2015)  *Scleroderma citrinum* (Kohler et al., 2015)  *Suillus luteus* (Kohler et al., 2015) |
| Secretomes from saprophytic fungi and yeasts | *Agaricus bisporus var burnettii* (Morin et al., 2012)  *Amanita thiersii* (Hess et al., 2014)  *Aspergillus niger* (Andersen et al., 2011)  *Aspergillus oryzae* (Machida et al., 2005)  *Coniophora puteana* (Floudas et al., 2012)  *Coprinopsis cinerea* (Stajich et al., 2010)  *Dacryopinax primogenitus* (Floudas et al., 2012)  *Dichomitus squalens* (Floudas et al., 2012)  *Fomitiporia mediterranea* (Floudas et al., 2012)  *Fomitopsis pinicola* (Floudas et al., 2012)  *Gloeophyllum trabeum* (Floudas et al., 2012)  *Gymnopus luxurians* (Kohler et al., 2015)  *Hydnomerulius pinastri* (Kohler et al., 2015)  *Hypholoma sublateritium* (Kohler et al., 2015)  *Neurospora crassa* (Galagan et al., 2003)  *Plicaturopsis crispa* (Kohler et al., 2015)  *Punctularia strigosozonata*  (Floudas et al., 2012)  *Sphaerobolus stellatus* (Kohler et al., 2015)  *Trametes versicolor* (Floudas et al., 2012)  *Wolfiporia cocos* (Floudas et al., 2012)  *Stereum hirsutum (*Floudas et al., 2012)  *Pseudozyma aphidis* (Lorenz et al., 2014)  *Pichia stipitis* (Jeffries et al., 2007)  *Pseudozyma antarctica* (Morita et al., 2013)  *Rhodosporidium toruloides* (Zhu et al., 2012)  *Saccharomyces cerevisiae* (Goffeau et al., 1996) |
| Secretomes from animal-pathogenic fungi | *Batrachochytrium dendrobatidis* (Rosenblum et al., 2010)  *Candida albicans* (Jones et al., 2004)  *Cordyceps militaris* (Zheng et al., 2011)  *Cryptococcus neoformans var. grubii* (Janbon et al., 2014)  *Cryptococcus neoformans var. neoformans* (Loftus et al., 2005)  *Encephalitozoon cuniculi* (Katinka et al., 2001)  *Enterocytozoon bieneusi* (Akiyoshi et al., 2009)  *Malassezia globosa* (Xu et al., 2007)  *Metarhizium robertsii* (Gao et al., 2011)  *Paracoccidioides brasiliensis* (Desjardins et al., 2011) |

**Supplementary Table S2: The machine learning models and training sets used for EffectorP 3.0.**

| **EffectorP 3.0 model** | **Positive training set** | **Negative training set** | **ML model** | **Included in EffectorP 3.0** |
| --- | --- | --- | --- | --- |
| Apoplastic model | Apoplastic fungal and oomycete effectors | Secreted proteins with an RxLR motif from oomycetes and secreted proteins from biotrophic/hemibiotrophic fungi | Naïve Bayes | Five best models |
|  |  |  | Decision tree | Five best models |
|  | Apoplastic fungal and oomycete effectors | Secreted proteins from fungal saprophytes and brown algae | Naïve Bayes | Five best models |
|  |  |  | Decision tree | Five best models |
|  | Apoplastic fungal and oomycete effectors | Secreted proteins from animal-pathogenic fungi and oomycetes | Naïve Bayes | Five best models |
|  |  |  | Decision tree | Five best models |
| Cytoplasmic model | Cytoplasmic fungal and oomycete effectors | Secreted oomycete proteins without an RxLR motif and secreted proteins from necrotrophic/hemibiotrophic fungi | Naïve Bayes | Five best models |
|  |  |  | Decision tree | Five best models |
|  | Cytoplasmic fungal and oomycete effectors | Secreted proteins from fungal saprophytes and brown algae | Naïve Bayes | Five best models |
|  |  |  | Decision tree | Five best models |
|  | Cytoplasmic fungal and oomycete effectors | Secreted fungal ectomycorrhizal proteins | Naïve Bayes | Five best models |
|  |  |  | Decision tree | Five best models |
| Cytoplasmic fungi-only model | Cytoplasmic fungal effectors | Secreted oomycete proteins without an RxLR motif and secreted proteins from necrotrophic/ hemibiotrophic fungi | Naïve Bayes | Five best models |
|  |  |  | Decision tree | Five best models |
|  | Cytoplasmic fungal effectors | Secreted proteins from fungal saprophytes and brown algae | Naïve Bayes | Five best models |
|  |  |  | Decision tree | Five best models |
|  | Cytoplasmic fungal effectors | Secreted fungal ectomycorrhizal proteins | Naïve Bayes | Five best models |
|  |  |  | Decision tree | Five best models |

**Supplementary Table S3: Independent validation set of effectors.**

| **Species** | **Effector** | **Effector localization** | **EffectorP 3.0 (probability)** | | **EffectorP-fungi 3.0 (probability)** | |
| --- | --- | --- | --- | --- | --- | --- |
|  |  |  | Cytoplasmic effector | Apoplastic effector | Cytoplasmic effector | Apoplastic effector |
| *Puccinia graminis* f. sp. *tritici* | AvrSr27 | Cytoplasmic | Y (0.722) | Y (0.772) | Y (0.871) | Y (0.772) |
| *Hemileia vastatrix* | AvrSh1 | Cytoplasmic | - | Y (0.606) | - | Y (0.606) |
| *Cladosporium fulvum* | Ecp11-1 | Apoplastic | Y (0.576) | Y (0.722) | - | Y (0.722) |
| *Blumeria graminis* | AvrPm1a | Cytoplasmic | Y (0.958) | - | Y (0.922) | - |
|  | AvrPm2 | Cytoplasmic | - | Y (0.59) | - | Y (0.59) |
|  | AVRa7 | Cytoplasmic | Y (0.73) | - | Y (0.596) | - |
|  | AVRa9 | Cytoplasmic | - | Y (0.887) | Y (0.66) | Y (0.887) |
|  | AVRa10 | Cytoplasmic | Y (0.692) | - | Y (0.836) | - |
| *Leptosphaeria maculans* | AvrLm6 | - | - | Y (0.73) | Y (0.512) | Y (0.73) |
| *Fusarium* *oxysporum* f. sp. *lycopersici* | SIX9 | - | Y (0.82) | Y (0.636) | - | Y (0.636) |
|  | SIX10 | - | - | Y (0.777) | - | Y (0.777) |
|  | SIX11 | - | Y (0.754) | Y (0.772) | - | Y (0.772) |
|  | SIX12 | - | - | Y (0.903) | - | Y (0.903) |
|  | SIX13 | - | Y (0.618) | - | - | - |
|  | SIX14 | - | - | Y (0.752) | - | Y (0.752) |
| *Magnaporthe oryzae* | MoCDIP1 | - | - | Y (0.563) | - | Y (0.563) |
|  | MoCDIP2 | - | - | Y (0.723) | - | Y (0.723) |
|  | MoCDIP3 | - | Y (0.729) | Y (0.865) | Y (0.824) | Y (0.865) |
|  | MoCDIP4 | - | - | Y (0.794) | - | Y (0.794) |
|  | MoCDIP5 | - | - | - | Y (0.508) | - |
|  | Avr-Pi54 | Cytoplasmic | - | - | Y (0.548) |  |
|  | Avr-Pita2 |  | Y (0.567) | Y (0.706) |  |  |
|  | Nis1 | Apoplastic | - | Y (0.587) |  |  |
| *Fusarium graminearum* | OSP24 | Cytoplasmic | Y (0.835) | Y (0.563) | Y (0.826) | Y (0.563) |
| *Ustilago maydis* | Mig2-1 | - | Y (0.738) | Y (0.824) | Y (0.858) | Y (0.824) |
|  | Mig2-3 | - | Y (0.556) | Y (0.79) | Y (0.57) | Y (0.79) |
|  | Mig2-4 | - | Y (0.753) | Y (0.648) | Y (0.824) | Y (0.648) |
|  | Mig2-5 | - | Y (0.723) | - | Y (0.633) | - |
|  | Mig2-6 | - | Y (0.611) | - | Y (0.689) | - |
|  | Hum3 | Apoplast | - | - | - | - |
|  | Rsp1 | Apoplast | Y (0.94) | - | Y (0.828) | - |
|  | Cce1 | Apoplast | Y (0.536) | Y (0.662) | Y (0.61) | Y (0.662) |
| *Hyaloperonospora arabidopsidis* | HaCR1 | Apoplast | Y (0.543) | Y (0.864) | Y (0.753) | Y (0.864) |
| *Plasmopara viticola* | PvRXLR53 | Cytoplasm | Y (0.788) | - | Y (0.704) | - |
| *Plasmopara viticola* | PvRXLR131 | Cytoplasm | Y (0.758) | - | Y (0.681) | - |
| *Phytophthora infestans* | PC2 | Apoplast | Y (0.785) | Y (0.888) | Y (0.699) | Y (0.888) |
|  | PexRD24 | Cytoplasm | Y (0.935) | - | Y (0.774) | - |
|  | EPIC3 | Apoplast | - | Y (0.849) | - | Y (0.849) |
|  | EPIC4 | Apoplast | - | - | - | - |
|  | EPI2 | Apoplast | Y (0.949) | Y (0.738) | Y (0.781) | Y (0.738) |
|  | EPI6 | Apoplast | Y (0.806) | - | Y (0.703) | - |
|  | SCR108 | Apoplast | Y (0.724) | Y (0.837) | Y (0.636) | Y (0.837) |
|  | PITG_22926 | Cytoplasm | Y (0.938) | - | Y (0.84) | - |
| *Phytophthora agathidicida* | PaRXLR24 | Cytoplasm | Y (0.923) | - |  |  |
| *Saprolegnia parasitica* | SpHTP1 | Cytoplasm | Y (0.76) | - | Y (0.734) | - |
|  | SpHTP3 | Cytoplasm | Y (0.789) | - | - | - |
| *Phytophthora sojae* | PSR2 | Cytoplasm | Y (0.766) | - | - | - |
| *Phytophthora parasitica* | AVH195 | Cytoplasm | Y (0.874) | - |  |  |

**Supplementary Table S4: Predicted cytoplasmic effectors in the baker’s yeast *Saccharomyces cerevisiae and their functional annotation.***

| **Yeast protein identifier** | **Annotation (UniProt)** | **Annotated localization (UniProt)** |
| --- | --- | --- |
| YGL228W | Outer spore wall assembly protein | Mitochondrion |
| YCR069W | Peptidyl-prolyl cis-trans isomerase | Membrane |
| YOL031C | Nucleotide exchange factor | Endoplasmic reticulum |
| YGR269W | Uncharacterized protein | - |
| YFR034W-A | Uncharacterized protein | - |
| YNL012W | Putative meiotic phospholipase | Nucleus, endoplasmic reticulum |
| YGR245C | Protein SDA1 | Nucleolus |
| YHR003C | tRNA threonylcarbamoyladenosine dehydratase 1 | Mitochondrion |
| YGL138C | Uncharacterized protein | - |
| YOR190W | Sporulation-specific glucan 1,3-beta-glucosidase | Secreted |
| YAR075W | Putative inosine-5'-monophosphate dehydrogenase 1 | Cytoplasm |
| YHR195W | Nucleus-vacuole junction protein 1 | Nucleus outer membrane |
| YCR101C | Uncharacterized protein | - |
| YPR016W-A | Uncharacterized protein | - |
| YGL020C | Golgi to ER traffic protein 1 | Membrane |
| YDR262W | FAS1 domain-containing protein | Vacuole |
| YNR066C | Uncharacterized membrane glycoprotein | Membrane |
| YBL008W-A | Uncharacterized protein | - |
| YML128C | Meiotic sister chromatid recombination protein 1 | - |
| YDR382W | 60S acidic ribosomal protein P2-beta | Cytoplasm |
| YFR039C | Outer spore wall protein 7 | - |
| YJR062C | Protein N-terminal amidase | - |
| YHR057C | Peptidyl-prolyl cis-trans isomerase B | Secreted |
| YML130C | Endoplasmic oxidoreductin-1 | Membrane |
| YPL123C | Ribonuclease T2-like | Vacuole, cytoplasm |
| YHR131W-A | Uncharacterized protein | - |
| YIR016W | Uncharacterized protein | - |
| YEL060C | Cerevisin | Vacuole |
| YNL185C | 54S ribosomal protein L19 | Mitochondrion |
| YNL190W | Hydrophilin | Cell wall |
| YLR385C | SWR1-complex protein 7 | Nucleus |
| YDR304C | Peptidyl-prolyl cis-trans isomerase D | Endoplasmic reticulum |
| YEL050C | 54S ribosomal protein RML2 | Mitochondrion |
| YBR096W | Uncharacterized protein | - |
| YOL068C | NAD-dependent protein deacetylase HST1 | Nucleus |
| YER170W | GTP:AMP phosphotransferase | Mitochondrion |
| YLR250W | Protein SSP120 | - |
| YPL011C | Transcription initiation factor TFIID subunit 3 | Nucleus |
| YJL192C | Suppressor of PMA1-7 protein 4 | Membrane |
| YOR288C | Protein disulfide-isomerase | Endoplasmic reticulum |
| YDR221W | Glucosidase 2 subunit beta | Endoplasmic reticulum |
| YMR065W | Nuclear fusion protein KAR5 | Membrane |
| YOR366W | Uncharacterized protein | - |
| YDR056C | Endoplasmic reticulum membrane protein complex subunit 10 | Membrane |
| YDR522C | Sporulation-specific protein 2 | Membrane |
| YKL021C | Maintenance of killer protein 11 | Membrane |
| YJL073W | DnaJ-like protein of the ER membrane 1 | Membrane |
| YNL093W | GTP-binding protein YPT53 | Membrane |

**Supplementary Table S5: Predicted apoplastic effectors in the baker’s yeast *Saccharomyces* *cerevisiae* and their functional annotation.**

| **Yeast protein identifier** | **Annotation (UniProt)** | **Annotated localization (UniProt)** |
| --- | --- | --- |
| YMR307W | 1,3-beta-glucanosyltransferase | GPI-anchor |
| YNL066W | Probable secreted beta-glucosidase | Cell wall |
| YKL223W | Putative UPF0377 protein | - |
| YBR013C | Uncharacterized protein | - |
| YHR126C | Probable GPI-anchored protein | GPI-anchor |
| YHR180W | Uncharacterized protein | Membrane |
| YBL108W | Putative UPF0377 protein | - |
| YKR042W | Probable secreted beta-glucosidase | Membrane |
| YKL164C | Cell wall mannoprotein | Cell wall |
| YOR382W | Facilitator of iron transport 2 | GPI-anchor |
| YPR016W-A | Uncharacterized protein | - |
| YOR072W | Uncharacterized protein | - |
| YHL037C | Uncharacterized protein | Secreted |
| YBL008W-A | Uncharacterized protein | Secreted |
| YGL258W | Velum formation protein 1 | Cytosol |
| YJL116C | Beta-glucosidase-like protein | Mitochondrial |
| YEL040W | Probable glycosidase CRH2 | GPI-anchor |
| YHR212W-A | Uncharacterized protein | - |
| YMR324C | Putative UPF0377 protein | Membrane |
| YBR096W | Uncharacterized protein | - |
| YER145C-A | Uncharacterized protein | - |
| YER188W | Uncharacterized protein | - |
| YML094C-A | Uncharacterized protein | - |
| YMR305C | Probable family 17 glucosidase SCW10 | Cell wall |
| YOL159C | Condition specific secretion protein 3 | Secreted |
| YOR387C | VEL1-related protein | Cytosol |
| YOL154W | Protein ZPS1 | - |
| YJR153W | Polygalacturonase | - |
| YJL160C | Cell wall protein PIR5 | Cell wall |
| YIR040C | Putative UPF0377 protein | - |
